# Age-dependent pathogenicity of two severe fever with thrombocytopenia syndrome viruses in a ferret model

**DOI:** 10.64898/2026.03.16.712105

**Authors:** Eun Bee Choi, Eun Young Jang, Seungyeon Kim, Seo Young Moon, Da-Young Kang, Hyo-Min Woo, Beoul Kim, You-Jeong Lee, Min-Goo Seo, Yoo-kyoung Lee, In-Ohk Ouh, Yong-Myung Kang

## Abstract

SFTSV is an emerging tick-borne pathogen associated with high case fatality rates, particularly in elderly patients. While severe pathogenicity has been reported in aged ferrets, lethal or clinically significant infection in younger animals and genotype-dependent differences in pathogenicity remain insufficiently defined. In this study, we established a ferret infection model using two Korean clinical isolates representing genotypes B and F and systematically compared disease progression between one-year-old and three-year-old ferrets. Three-year-old ferrets exhibited rapid fever onset, marked body weight loss, early clinical deterioration, severe thrombocytopenia and leukopenia, significant elevations in AST and ALT levels, and earlier peak viremia with higher tissue viral loads, indicating impaired early viral control and accelerated systemic dissemination. Notably, one-year-old ferrets also developed measurable pathogenic manifestations, including febrile responses, progressive weight loss, detectable viremia, and multiorgan viral distribution, although disease progression was delayed and less severe compared with older animals. Within the same age group, differences in pathogenicity between genotypes B and F were limited. These findings demonstrate that host age is a major determinant of SFTSV disease severity and support the use of an age-stratified ferret model for preclinical evaluation of vaccines and antiviral therapeutics.

**Importance:** SFTS is an emerging tick-borne disease that can cause high fever, thrombocytopenia, and multi-organ failure, with particularly severe outcomes in older adults. Currently, no approved vaccines or specific antiviral treatments are available. Reliable animal models that recapitulate human disease are therefore essential for the development of effective countermeasures. Ferrets have recently been proposed as a useful model for SFTS, especially in aged animals, but the susceptibility of younger ferrets and the impact of viral strain differences remain unclear. Here we show that host age strongly determines disease severity in ferrets infected with two genetically distinct SFTS virus strains, establishing a flexible animal model for evaluating vaccines and antiviral therapies.

## Introduction

Severe fever with thrombocytopenia syndrome (SFTS) is an emerging tick-borne viral disease associated with high fever, thrombocytopenia, leukopenia, and multiorgan dysfunction (1). Since its emergence in East Asia, SFTS has posed a significant public health concern due to its relatively high case fatality rate and the absence of approved vaccines or specific antiviral therapies (2–4). Clinical epidemiological data consistently indicate that disease severity and mortality are markedly higher in elderly patients, highlighting age as a critical risk factor for severe outcomes(5, 6). These characteristics underscore the urgent need for effective preventive strategies, particularly vaccines capable of protecting high-risk populations.

The development of effective vaccines and therapeutics for SFTS depends on animal models that accurately recapitulate key features of human disease, including fever, weight loss, hematological abnormalities, hepatic injury, viremia, and systemic viral dissemination, as well as age-dependent differences in disease severity (7). However, currently available models each have important limitations: immunodeficient mouse models can develop severe disease but are unsuitable for vaccine evaluation, while immunocompetent rodents, hamsters, and non-human primates generally exhibit mild or non-lethal infection and fail to reproduce severe human SFTS (8–10). Consequently, there remains a critical need for an alternative animal model that retain intact immune function while exhibiting clinically relevant, age-sensitive disease manifestations.

Ferrets (Mustela putorius furo) are well-established models for human viral infections and share physiological and immunological similarities with humans (11, 12). Recent studies have demonstrated that ferrets infected with SFTSV develop clinical signs and laboratory abnormalities resembling those seen in human SFTS patients, including fever, weight loss, thrombocytopenia, and leukopenia (13). Notably, aged ferrets exhibit more severe disease outcomes than younger animals, suggesting that ferrets may provide a valuable platform for studying age-dependent SFTSV pathogenesis (13, 14). However, systematic comparisons incorporating host age and viral diversity under standardized experimental conditions remain limited, particularly for SFTSV isolates circulating in Korea.

In this study, we studied the age-dependent pathogenicity of SFTSV using a ferret infection model. One-year-old and three-year-old ferrets were inoculated with two genetically distinct SFTSV isolates obtained in Korea, and disease progression was evaluated using comprehensive clinical, hematological, biochemical, and virological parameters. By directly comparing disease outcomes across age groups and viral isolates, this study aims to establish a standardized, age-sensitive ferret model of SFTSV infection and to provide a robust preclinical platform for the evaluation of candidate vaccines and therapeutics against SFTS.

## Results

### Age-dependent changes in body temperature and body weight following SFTSV infection

To evaluate age-dependent differences in disease severity following SFTSV infection, changes in body temperature and body weight were monitored daily in one-year-old and three-year-old ferrets inoculated with Korean isolates of SFTSV B type or F type. PBS-inoculated control animals in both age groups maintained stable body temperature and body weight throughout the observation period, indicating that the experimental procedures themselves did not induce detectable systemic stress or illness (Fig 1).

**Fig 1.**
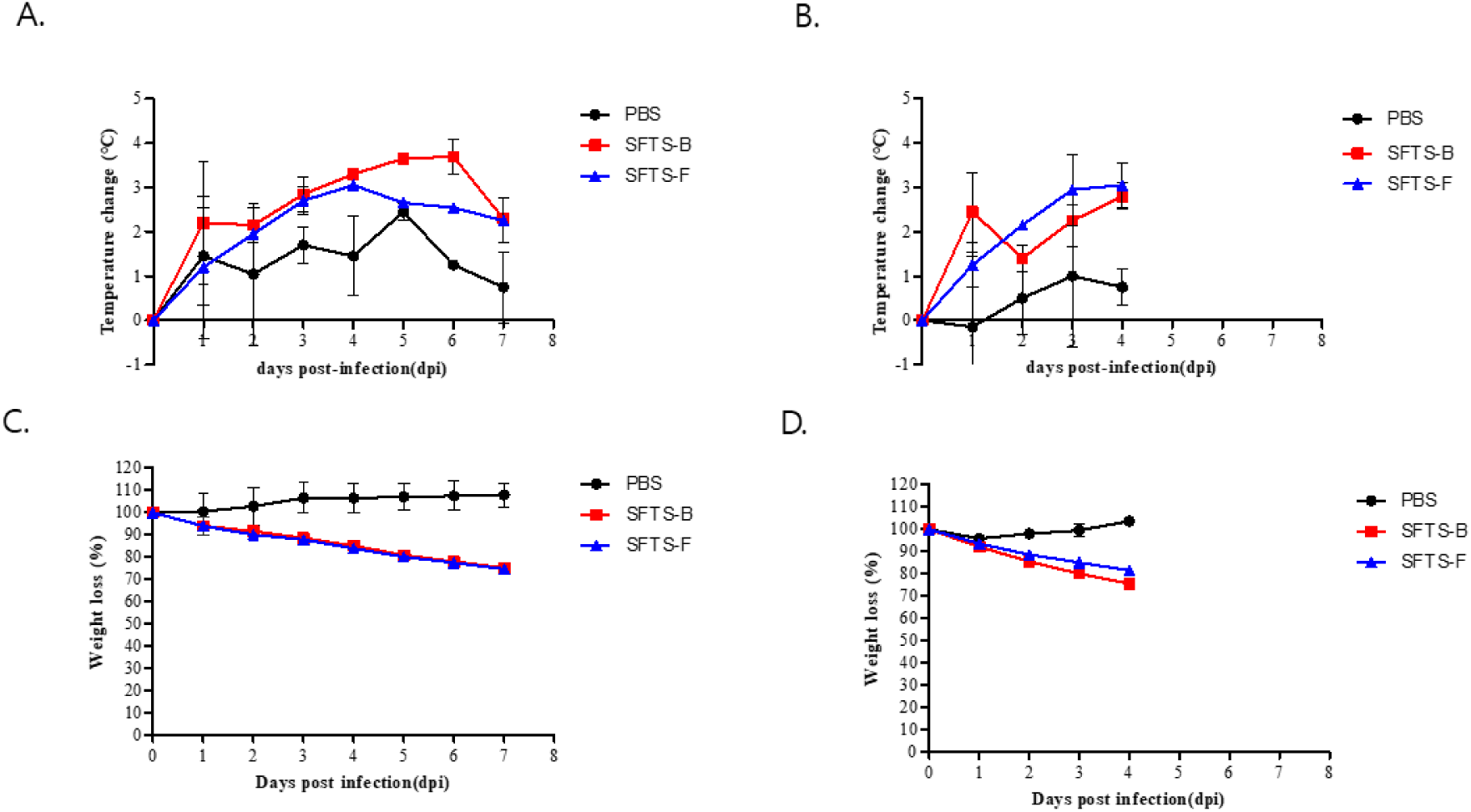
Age-dependent clinical manifestations following SFTSV infection in Ferrets. **C**hanges in body temperature and body weight were monitored in one-year-old and three-year-old ferrets following infection with SFTSV genotype B or F. (A) Temperature changes in one-year-old ferrets. (B) Temperature changes in three-year-old ferrets. (C) Percentage of body weight change in one-year-old ferrets. (D) Percentage of body weight change in three-year-old ferrets. Three-year-old ferrets exhibited earlier onset of fever and greater body weight loss compared with one-year-old animals. Data are shown as mean ± SD (n = 2 per group). PBS, phosphate-buffered saline control.

In SFTSV-infected ferrets, increases in body temperature accompanied by progressive body weight loss were observed in both age groups. In one-year-old ferrets, body temperature increased from 1 day post infection (dpi) and remained elevated between approximately 3 and 6 dpi in both viral groups. Body weight loss became apparent from 1 dpi and progressed gradually, reaching up to approximately 20% by 7 dpi. Within this age group, no consistent differences in the patterns of temperature change or body weight loss were observed between animals infected with SFTSV B type and those infected with F type (Fig 1A and 1C).

In three-year-old ferrets, increases in body temperature and body weight loss were observed earlier after infection. Elevated body temperature was detected from 1 dpi, and body weight declined rapidly, with animals reaching the predefined humane endpoint ( ≥ 20% body weight loss) by 4 dpi, at which point euthanasia was required. Although formal statistical comparisons were not performed due to the limited number of animals per group, the earlier onset of fever and accelerated weight loss observed in older ferrets suggest increased susceptibility to SFTSV-associated systemic disease in this age group (Fig 1B and 1D).

### Clinical symptom scores following SFTSV infection

To further characterize clinical disease progression, ferrets were evaluated daily using a standardized clinical scoring system. PBS-inoculated control animals in both age group remained asymptomatic throughout the observation period (Fig 2). In one-year-old ferrets infected with SFTSV, clinical signs such as reduced activity, lethargy, and decreased appetite became apparent from 3 dpi and gradually increased in severity overtime. Animals infected with SFTSV B type and F type exhibited comparable trajectories of clinical score increases, with peak scores observed between 6 and 7 dpi (Fig 2A).

**Fig 2.**
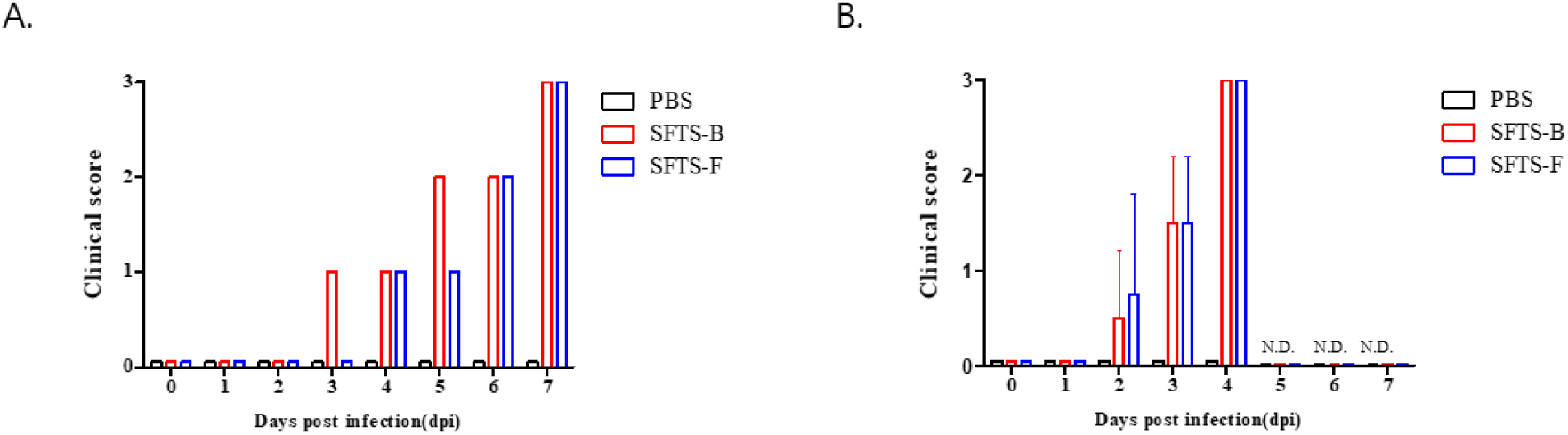
Clinical score progression in SFTSV-infected ferrets. Clinical scores were assessed daily based on activity level, posture, and appetite. (A) Clinical scores in one-year-old ferrets. (B) Clinical scores in three-year-old ferrets. Older ferrets showed more rapid progression of clinical deterioration. N.D., not determined due to humane endpoint (n = 2 per group).

In three-year-old ferrets, clinical signs appeared earlier after infection and progressed more rapidly. Increases in clinical scores were observed from 2 dpi, with scores rising sharply and reaching peak levels by 3–4 dpi, prior to animals meeting predefined humane endpoints. Due to the limited number of animals per group, formal statistical comparisons were not performed; however, the earlier onset and accelerated progression of clinical signs observed in older ferrets are consistent with increased disease severity in this age group (Fig 2B)

### Age-dependent hematological abnormalities and liver injury following SFTSV infection

To assess systemic inflammation and organ injury associated with SFTSV infection, hematological parameters and serum biochemical markers were analyzed in one-year-old and three-year-old ferrets inoculated with SFTSV B type or F type. PBS-inoculated control animals in both age groups maintained stable white blood cell (WBC) counts, platelet (PLT) levels, and serum aspartate aminotransferase (AST) and alanine aminotransferase (ALT) concentrations throughout the experimental period (Fig 3 & 4).

**Fig 3.**
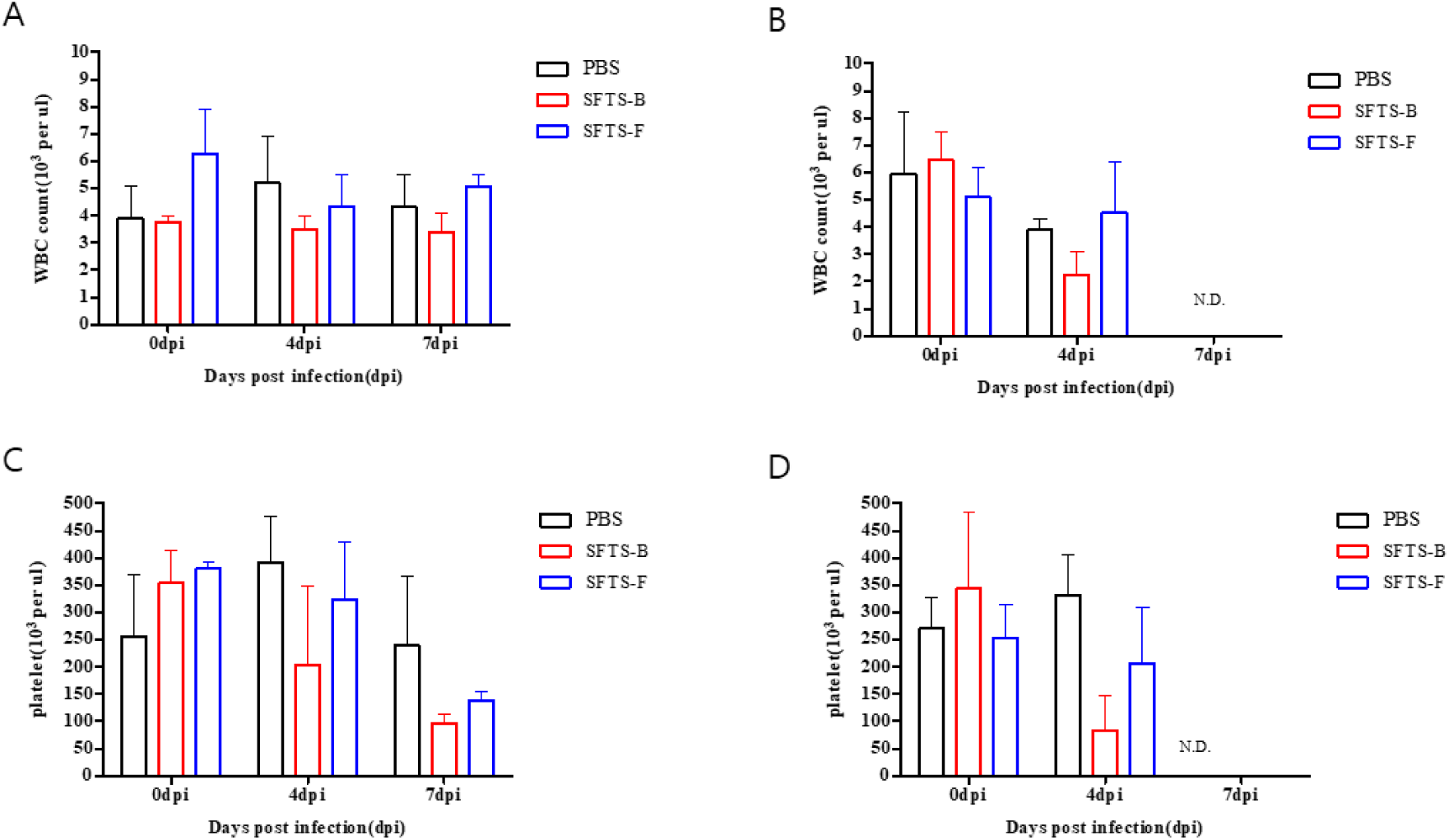
Hematological alterations following SFTSV infection. White blood cell (WBC) and platelet (PLT) counts were measured at 0, 4, and 7 days post-infection (dpi). (A) WBC counts in one-year-old ferrets. (B) WBC counts in three-year-old ferrets. (C) Platelet counts in one-year-old ferrets. (D) Platelet counts in three-year-old ferrets. Three-year-old ferrets developed rapid leukopenia and severe thrombocytopenia compared with younger animals. Data represent mean ± SD (n = 2 per group). N.D., not determined.

**Fig 4.**
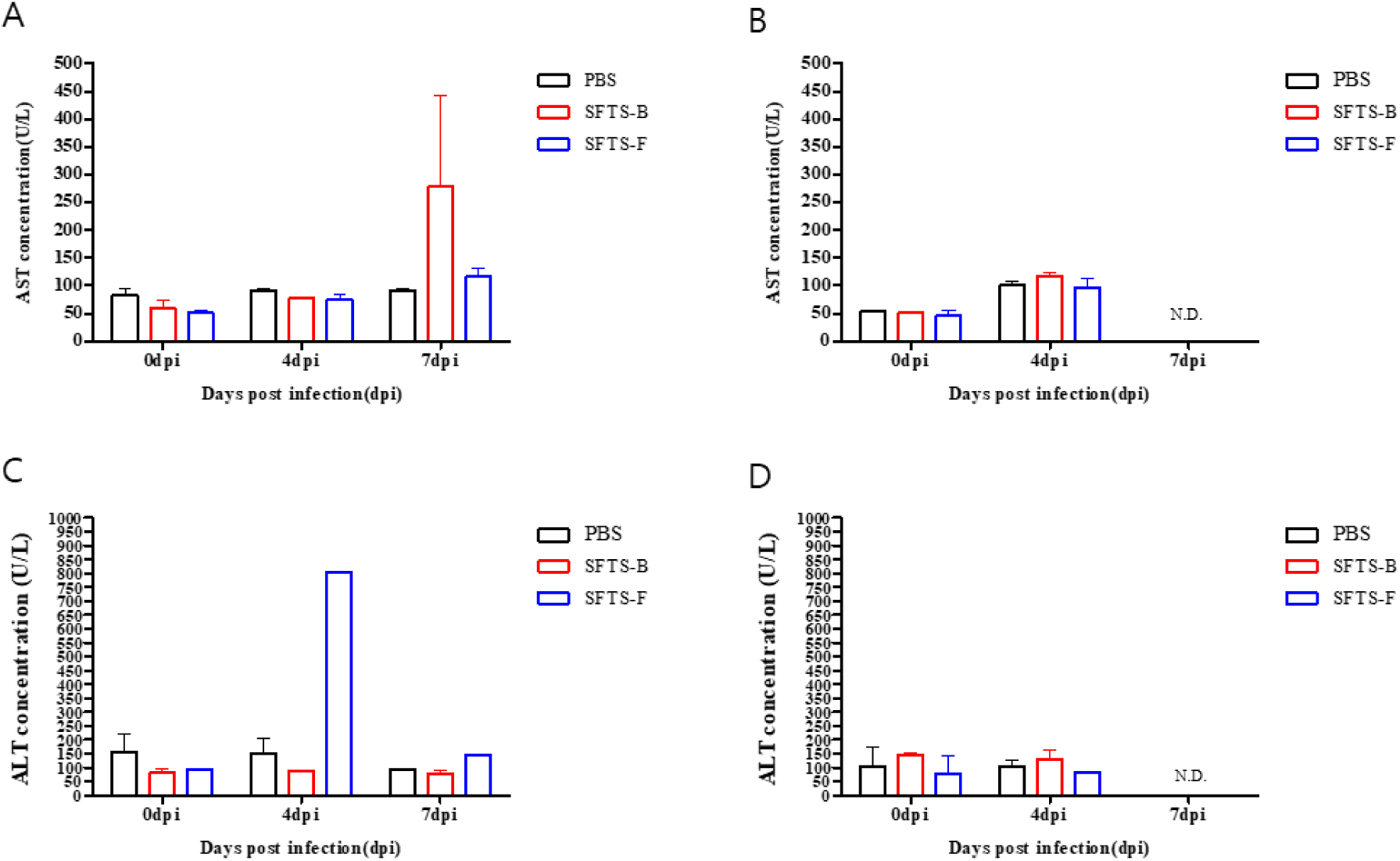
Serum biochemical parameters following SFTSV infection. Serum aspartate aminotransferase (AST) and alanine aminotransferase (ALT) levels were evaluated at 0, 4, and 7 dpi. (A) AST levels in one-year-old ferrets. (B) AST levels in three-year-old ferrets. (C) ALT levels in one-year-old ferrets. (D) ALT levels in three-year-old ferrets. Marked elevations in AST and ALT were observed in infected animals, particularly in three-year-old ferrets, indicating hepatic injury. Data are shown as mean ± SD (n = 2 per group). N.D., not determined.

In one-year-old ferrets, SFTSV infection was associated with moderate hematological and biochemical changes. WBC counts generally remained within the normal range over the course of infection, whereas platelet counts showed a gradual decline, becoming most evident at 7 dpi in both viral groups (Fig 3A & 3C). In parallel, serum AST levels increased progressively, suggesting developing hepatic involvement (Fig 4A), while ALT levels showed a milder elevation (Fig 4C). An increase in ALT was particularly noticeable at 4 dpi in ferrets infected with SFTSV F type. Collectively, these observations indicate the development of subacute systemic inflammation and liver injury in younger ferrets following SFTSV. In three-year-old ferrets, hematological and biochemical abnormalities appeared earlier and progressed more rapidly. Reduction in platelet and WBC counts were observed by 4 dpi in animals infected with either SFTSV B type or F type (Fig 3B & 3D). At the same time, serum AST and ALT levels increased markedly, consistent with acute liver injury (Fig 4B & 4D).

### Age-dependent virus shedding by rectal swab following SFTSV infection

To assess viral shedding following SFTSV infection, rectal swab samples were collected daily and viral loads were quantified as FFU/mL equivalents (Fig 5). In the one-year-old ferret group, detectable viral shedding was observed as early as 1 dpi in some animals and persisted intermittently through the experimental period, with peak viral loads generally observed between 1 and 2 dpi. Both viral genotypes exhibited comparable shedding patterns, although the magnitude and timing of viral detection varied among individual animals (Fig 5A). In the three-year-old ferret group, rectal swab viral detection was more limited. Viral RNA was detected primarily at 3–4 dpi in both SFTSV B type– and F type–infected animals. Overall, viral shedding in rectal swabs was transient in older ferrets and appeared to coincide with the acute phase of systemic infection (Fig 5B).

**Fig 5.**
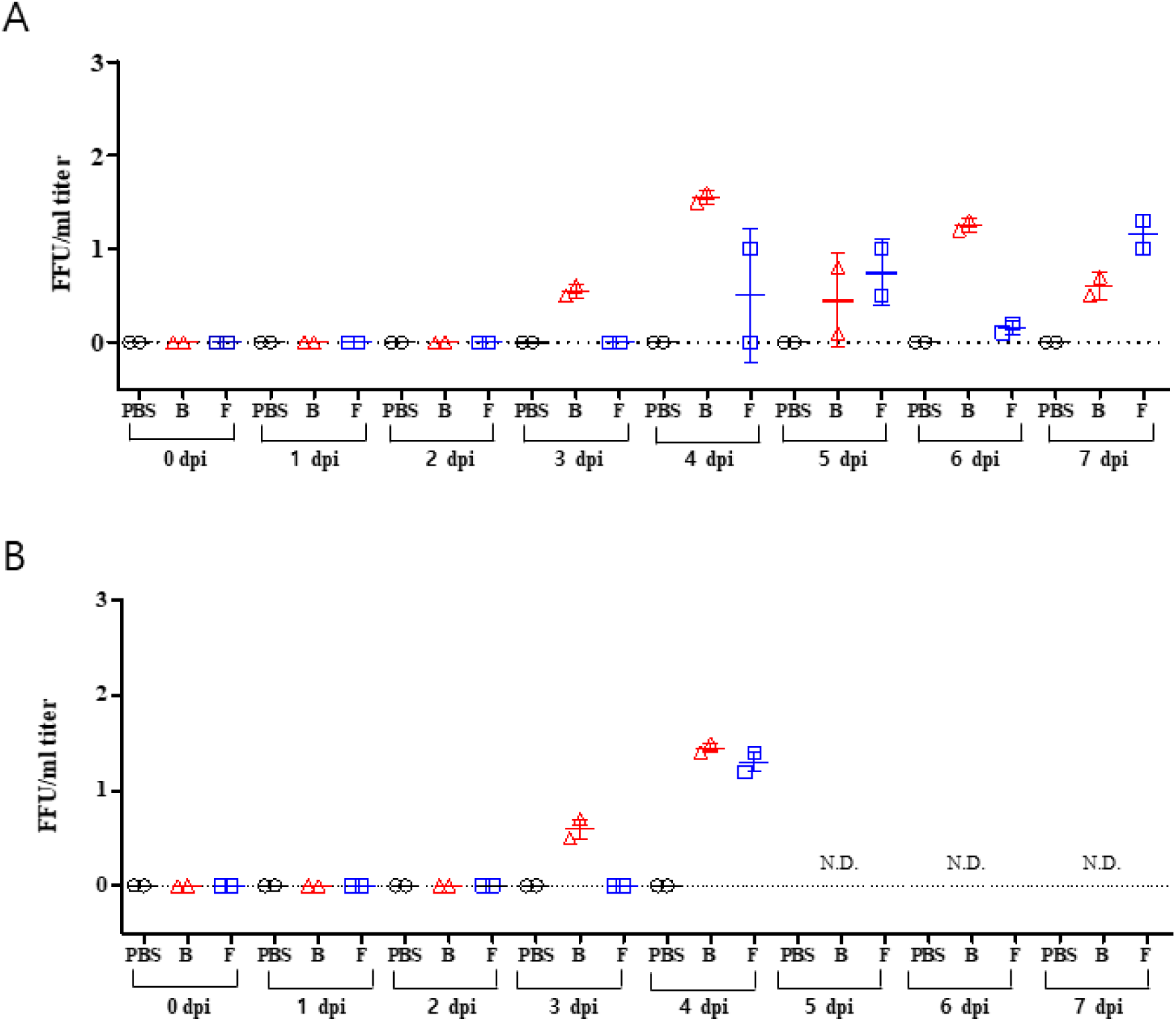
Serum viral titers following SFTSV infection. Serum viral loads were determined by real-time RT-PCR and converted to FFU/mL equivalents using a standard curve generated from virus stocks with known infectious titers. (A) Viral titers in one-year-old ferrets from 0 to 7 dpi. (B) Viral titers in three-year-old ferrets. Older ferrets exhibited earlier peak viremia compared with younger animals. Each symbol represents an individual animal (n = 2 per group). N.D., not determined.

No viral RNA was detected in rectal swab samples collected from PBS-inoculated control animals at any time point. Collectively, these findings indicate that SFTSV infection results in detectable viral shedding via the gastrointestinal tract in both age groups, with more prolonged and variable shedding observed in one-year-old ferrets.

### Age-dependent systemic viral dissemination and tissue tropism following SFTSV infection

To investigate tissue tropism and systemic dissemination of SFTSV, viral loads were quantified in major organs and blood at necropsy (Fig 6). In one-year-old ferrets, SFTSV was detected in multiple organs following infection with both B type and F type viruses. Among the tissues examined, the spleen consistently exhibited the highest viral loads, followed by the liver and small intestine, indicating preferential viral replication in lymphoid and visceral organs. Moderate viral loads were also detected in the kidney, whereas viral detection in the brain was absent. Low-level viremia was observed in blood samples, consistent with systemic viral spread (Fig 6A).

**Fig 6.**
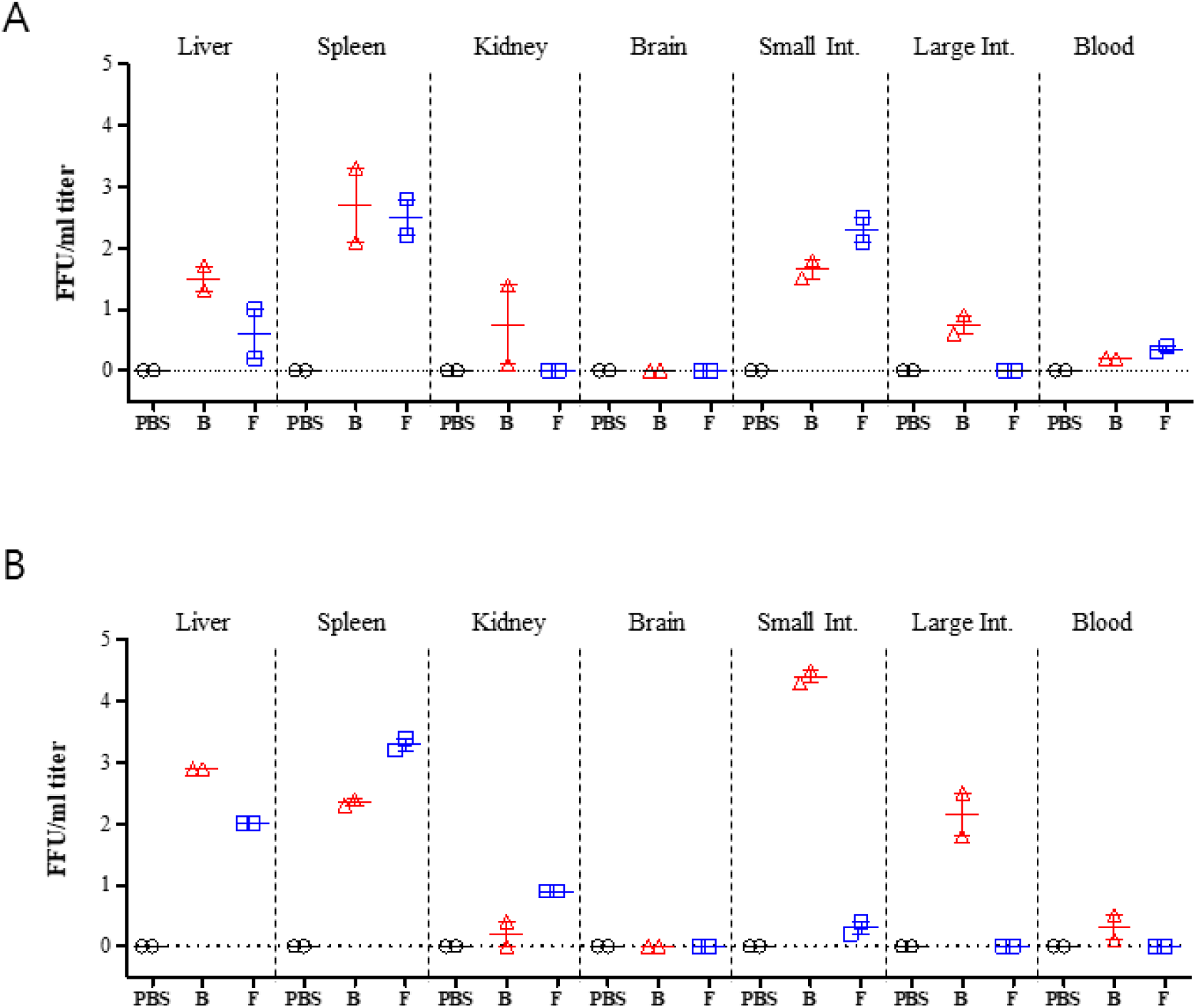
Tissue viral distribution in major organs. Viral titers in liver, spleen, kidney, brain, small intestine, large intestine, and blood were quantified at necropsy and expressed as FFU/mL equivalents. (A) One-year-old ferrets. (B) Three-year-old ferrets. The spleen consistently exhibited the highest viral burden across groups, followed by liver and small intestine. Viral detection in the brain was minimal. Each symbol represents an individual animal (n = 2 per group).

In three-year-old ferrets, viral dissemination was more pronounced and showed a similar but amplified tissue distribution pattern. High viral loads were detected in the spleen and liver in both SFTSV B type– and F type–infected animals. Notably, the small intestine exhibited particularly high viral loads in the B type–infected group, suggesting substantial gastrointestinal involvement in older animals. Viral loads in the kidney were detectable but lower than those observed in the spleen and liver, while viral detection in the brain remained limited. Blood samples from infected animals contained detectable viral loads, reflecting acute systemic viremia prior to euthanasia.

Across both age groups, PBS-inoculated control animals showed no detectable viral signal in any tissue or blood sample. Overall, these findings demonstrate that SFTSV infection results in broad tissue dissemination with a marked tropism for the spleen, liver, and gastrointestinal tract. Although both viral genotypes exhibited similar tissue distribution patterns, older ferrets tended to show higher viral burdens across multiple organs, consistent with the more severe clinical course observed in this age group.

### Histopathological changes in the spleen and liver following SFTSV infection

Histopathological examination revealed distinct inflammatory changes in the spleen and liver of SFTSV-infected ferrets compared with control animals. In the spleen, the PBS control groups (Fig. 7A, 7D) exhibited normal splenic architecture, with intact and clearly demarcated boundaries between the white pulp (w) and red pulp (r), and no remarkable histopathological lesions. In contrast, SFTSV-infected animals showed distinct inflammatory and degenerative alterations in the spleen. In both the B type and F type groups, there was a diffuse expansion/accumulation of large macrophage-like cells throughout the splenic parenchyma (Fig. 7B, 7C, 7E, 7F). This change was most evident within the red pulp and is highlighted by open arrows, with the F type-infected animals demonstrating a more prominent histiocytic response, particularly in Fig. 7C and 7F. Additionally, lymphocytic necrosis within the white pulp was observed to varying degrees in both infection groups. These lesions, associated with lymphoid depletion and disruption of the white pulp architecture (indicated by small black arrows), were generally more pronounced in the B type group (Fig. 7B, 7E). Collectively, these findings indicate that SFTSV infection induces substantial splenic pathology, characterized by prominent macrophage-like cell expansion in the F type group and comparatively more severe white pulp lymphocytic necrosis in the B type group.

**Fig. 7.**
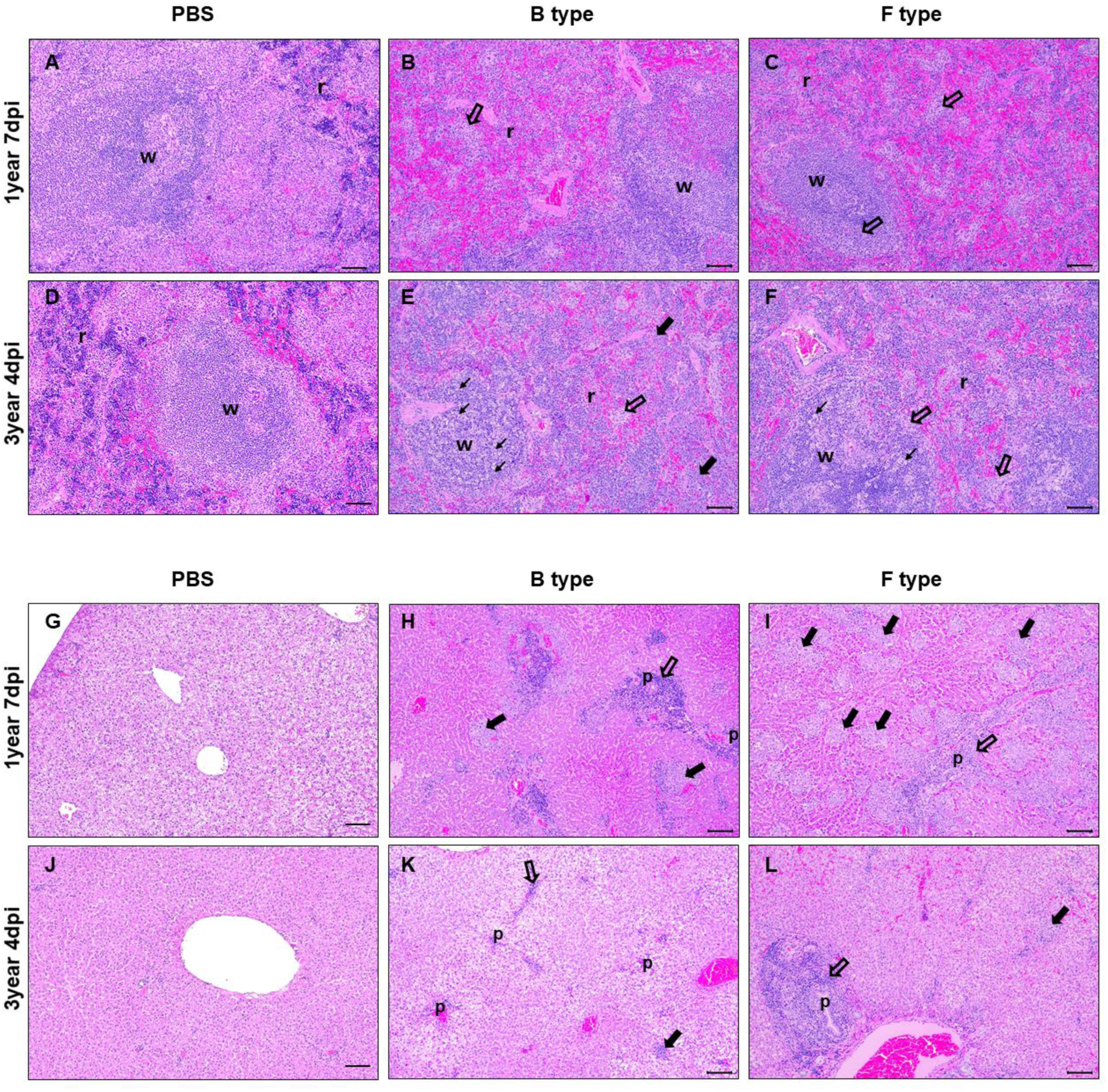
Histopathological changes in the spleen and liver following SFTSV infection. Representative hematoxylin and eosin (H&E)-stained sections of spleen (A–F) and liver (G–L) from PBS-treated controls and SFTSV-infected ferrets. (A) Spleen from a one-year-old PBS-treated ferret showing normal splenic architecture with well-demarcated white pulp (w) and red pulp (r). (B, C) Spleens from one-year-old ferrets infected with SFTSV B type (B) or F type (C) showing prominent red pulp expansion with diffuse accumulation/expansion of large macrophage-like cells (open arrows) and relative attenuation/partial depletion of the white pulp (w). (D) Spleen from a three-year-old PBS-treated ferret showing preserved white pulp–red pulp architecture. (E, F) Spleens from three-year-old ferrets infected with SFTSV B type (E) or F type (F) demonstrating white pulp lymphoid injury with lymphoid depletion and necrotic changes within lymphoid follicles (small black arrows), accompanied by increased red pulp histiocytic expansion (open arrows). r, red pulp; w, white pulp. (G) Liver from a one-year-old PBS-treated ferret showing normal hepatic architecture. (H, I) Liver sections from one-year-old ferrets infected with SFTSV B type (H) or F type (I) showing multifocal mononuclear inflammatory foci within hepatic lobules and periportal/portal inflammatory cell infiltration (p; open arrows), with scattered foci of hepatocellular degeneration and/or necrosis (solid black arrows). (J) Liver from a three-year-old PBS-treated ferret showing preserved lobular organization. (K, L) Liver sections from three-year-old ferrets infected with SFTSV B type (K) or F type (L) showing persistent and more prominent portal and periportal mononuclear cell infiltration (p; open arrows), lobular inflammatory nodules (solid black arrows), and hepatocellular degeneration with variable vacuole accumulation. p, portal area. Scale bars as indicated.

To assess the extent of hepatic injury following SFTSV infection, a histopathological evaluation was performed on liver tissues from PBS-treated controls, B type-infected, and F type-infected groups (Fig. 7G–7L). The PBS control groups (Fig. 7G, 7J) exhibited normal hepatic architecture with preserved lobular organization and intact hepatocellular cords, and no appreciable inflammatory or necrotic lesions. In contrast, prominent pathological alterations were observed in infected animals. Both B type and F type groups showed multifocal inflammatory foci within the hepatic lobules, predominantly composed of mononuclear cells, as well as similar inflammatory cell infiltration in the portal tracts (p) and surrounding periportal regions (open arrows), consistent with portal/periportal hepatitis (Fig. 7H, 7I, 7K, 7L). These inflammatory changes were accompanied by scattered foci of hepatocellular injury, including degeneration and/or spotty to focal necrosis (solid black arrows), indicating concurrent parenchymal damage. The severity and distribution of lesions differed between the infection groups. F type-infected animals (Fig. 7I, 7L) exhibited more extensive and multifocal hepatocellular necrosis and degeneration at 1-year 7 dpi, and subsequently showed severe, widespread periportal inflammation at 3-year 4 dpi with partial disruption of the surrounding tissue architecture. Although the B type group (Fig. 7H, 7K) also demonstrated portal/periportal mononuclear inflammation and focal hepatocellular necrosis, the overall extent and severity were comparatively less pronounced than those observed in the F type group. In addition, varying degrees of hepatocellular vacuole accumulation were noted in infected animals, suggesting concurrent hepatic injury and metabolic alteration (Fig. 7G–7L). Collectively, these findings indicate that SFTSV infection induces significant hepatic pathology characterized by lobular and portal-based mononuclear inflammation with accompanying hepatocellular degeneration/necrosis and lipid accumulation, with the F type infection producing a more severe and extensive hepatic response than the B type infection.

## Discussion

In this study, we established a ferret model of SFTSV infection using two Korean clinical isolates representing genotypes B and F, and systematically evaluated age-dependent differences in pathogenicity, viral dissemination, and tissue tropism to infer their relevance to human disease. By comparing one-year-old and three-year-old ferrets under identical experimental conditions, we demonstrated that host age is a critical determinant of disease severity, while viral genotype also contributed to differences in clinical manifestations and pathogenic outcomes.

One of the most notable findings of this study is the pronounced age-dependent divergence in disease progression following SFTSV infection. Compared with one-year-old animals, three-year-old ferrets exhibited more rapid onset of fever, accelerated body weight loss, earlier deterioration of clinical signs, and mortality. Analyses of serum viremia and tissue viral distribution further revealed that older ferrets reached peak viremia earlier and harbored higher viral loads in multiple organs prior to death, indicating impaired early viral control and rapid systemic dissemination. These features closely resemble the clinical course of severe SFTS in elderly human patients, in whom advanced age is a well-established risk factor for disease severity and fatal outcome (15, 16). The heightened pathogenicity observed in older ferrets may reflect age-associated alterations in antiviral immune responses, dysregulated inflammatory control, or reduced tissue resilience, all of which could increase susceptibility to SFTSV infection.

Importantly, our results demonstrate that one-year-old ferrets also developed clear pathogenic manifestations following SFTSV infection in contrast to several previous ferret studies. Earlier reports have largely described overt disease only in aged ferrets, typically ≥ 4 years of age, while younger animals were considered asymptomatic or minimally affected (13, 17, 18). In the present study, however, one-year-old ferrets consistently exhibited febrile responses, progressive body weight loss, increased clinical scores, detectable viremia, and viral dissemination to multiple organs. Although disease progression in younger ferrets was slower and less severe than in three-year-old animals, the presence of clear clinical, hematological, and virological abnormalities relative to control animals indicates an intermediate pathogenic phenotype rather than complete resistance. This finding expands the current understanding of age-dependent susceptibility in the ferret model and suggests that SFTSV pathogenicity occurs along a continuum rather than being restricted to a narrow age threshold. Several factors may account for the discrepancy between our findings and those of earlier studies. The use of recent Korean clinical isolates with potentially higher intrinsic pathogenicity may have contributed to enhanced disease expression. In addition, the relatively high inoculation dose and the use of multiple inoculation routes may have facilitated systemic viral dissemination even in younger animals. Together, these experimental conditions may better approximate severe natural exposure scenarios and reveal pathogenic features that were not apparent under more restrictive or lower-dose challenge models.

Consistent with clinical observations, age-dependent differences were also evident in hematological and biochemical parameters. Three-year-old ferrets developed rapid and severe thrombocytopenia and leukopenia, accompanied by early and marked elevations in serum AST and ALT levels, indicative of acute systemic inflammation and hepatic injury. These findings are in line with the typical laboratory features reported in severe human SFTS (19–21). In contrast, one-year-old ferrets showed more gradual hematological changes and delayed liver enzyme elevations, consistent with a subacute disease course. Within the same age group, however, differences between SFTSV genotypes B and F were relatively limited. Fever kinetics, weight loss, clinical scores, hematological indices, biochemical markers, and viral loads were largely comparable between genotypes, suggesting that under high-dose experimental conditions, host factors—particularly age—exert a stronger influence on disease outcome than viral genotype (17).

Analysis of tissue viral distribution revealed consistent organ tropism across experimental groups. The spleen harbored the highest viral loads, highlighting its role as a major target organ for SFTSV replication. Substantial viral burdens were also detected in the liver and small intestine, indicating extensive visceral involvement during systemic infection (13, 22). In contrast, viral detection in the brain was minimal, suggesting limited neurotropism under the experimental conditions employed. Detection of viral RNA in rectal swab samples further indicates active viral shedding following systemic infection. Although tick-borne transmission is the primary route for SFTSV, gastrointestinal shedding may contribute to environmental contamination or indirect exposure risks in severe cases, underscoring the value of non-invasive sampling strategies in preclinical studies (22).

Several limitations of this study should be acknowledged. The relatively small group sizes may limit the statistical power to detect subtle genotype-dependent differences. In addition, immunological correlates of disease severity or protection were not directly assessed. Future studies incorporating cytokine profiling, neutralizing antibody responses, and cellular immune analyses will be essential to elucidate the mechanisms underlying age-dependent susceptibility and disease progression.

In conclusion, our findings demonstrate that SFTSV infection in ferrets is strongly influenced by host age, with older animals exhibiting accelerated disease progression, enhanced viral dissemination, and increased multiorgan involvement. Importantly, this study also shows that pathogenic SFTSV infection can occur in younger ferrets, challenging the prevailing view that overt disease is restricted to aged animals. Together, these results support the establishment of a flexible and clinically relevant ferret model that can accommodate a spectrum of disease severity and provide a robust platform for the preclinical evaluation of vaccines and antiviral therapeutics against SFTS.

## Materials and methods

### Viruses and cells

Two genetically distinct isolates of SFTSV circulating in Korea were used in this study. The viruses were classified as SFTSV B type (GenBank accession numbers KP663743–KP663745), isolated from South Korea in 2014, and SFTSV F type (GenBank accession numbers KF358691–KF358693), isolated from South Korea in 2012, based on phylogenetic analysis of viral genome sequences.

Viral stocks were propagated in Vero cells (ATCC no.CRL-1586; American Type Culture Collection), which were maintained in Minimum Essential Medium (MEM; Gibco, USA) supplemented with 2% FBS (Gibco), 1% penicillin–streptomycin, and incubated at 37 °C with 5% CO2. Virus-containing supernatants were harvested when cytopathic effects became evident, clarified by low-speed centrifugation, aliquoted, and stored at −80 °C until use.

Viral titers were determined by focus-forming assay (FFA) assay on Vero cells and expressed as FFU/mL. Prior to animal inoculation, viral titers were reconfirmed by back titration to ensure accurate dosing.

### Ferret infection study

Specific-pathogen-free (SPF) female ferrets (Mustela putorius furo) were obtained and divided into two age groups: one-year-old (young adult) and three-year-old (adult) animals. Ferrets were housed individually in biosafety level 3 (BSL-3) animal facilities with controlled temperature and humidity and provided food and water ad libitum.

Each experimental group consisted of two ferrets. Ferrets were inoculated with either SFTSV B type or SFTSV F type virus at a concentration of 10^7^ FFU/mL. Each animal received a total inoculation volume of 4 mL, which was administered via a combination route to facilitate systemic injection : 1mL by subcutaneous (SC), 1mL by intramuscular (IM), and 2mL by intraperitoneal (IP) routes. Control animals were inoculated with an equivalent volume and route of PBS. Animals were monitored daily for clinical signs, including activity level, posture, appetite, and survival [Table 1].

**Table 1.**
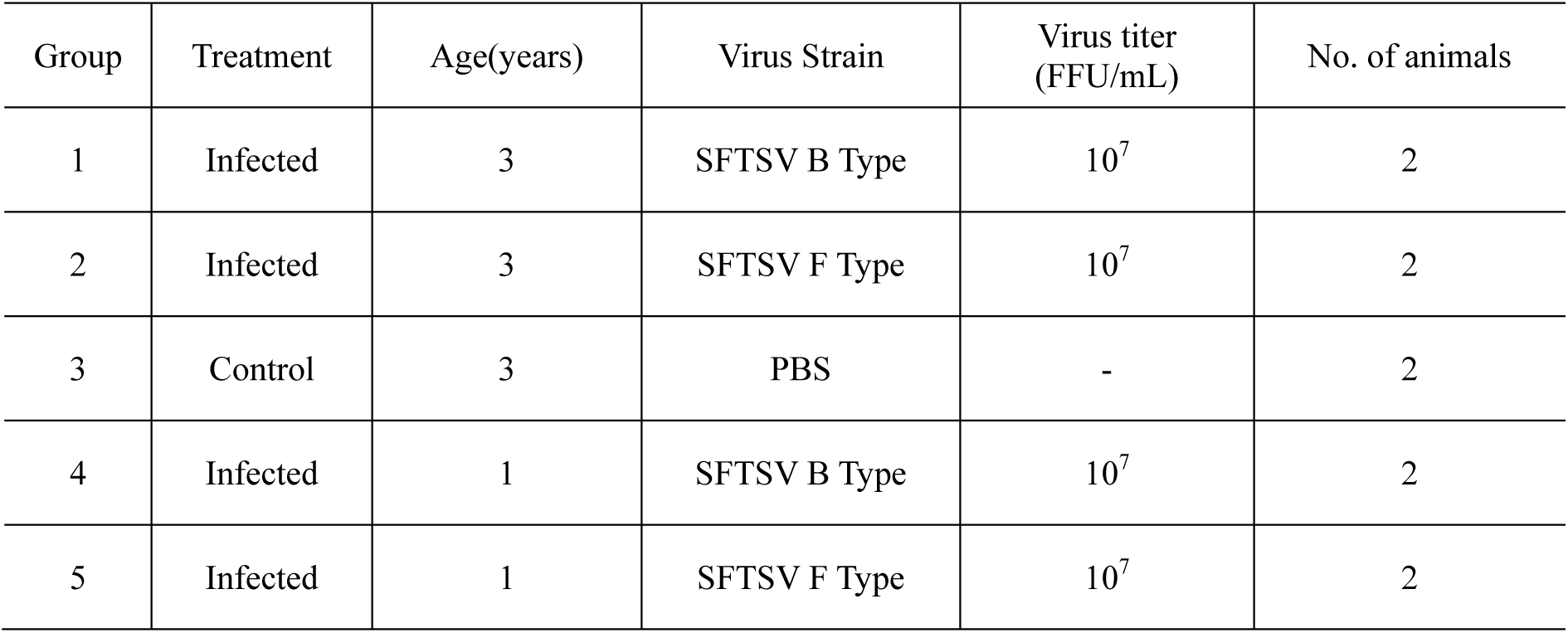
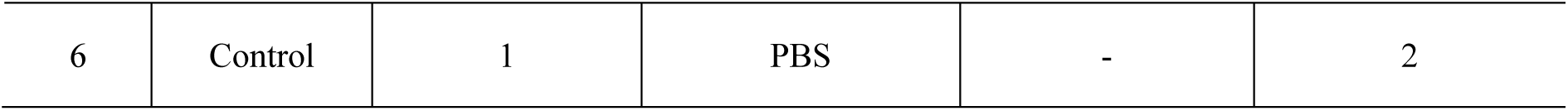
Experimental design of the SFTSV infection study in Ferrets. Ferrets were inoculated with SFTSV genotype B or F at the indicated titers. Each animal received a total inoculation volume of 4 mL administered via combined routes (1 mL subcutaneous, 1 mL intramuscular, and 2 mL intraperitoneal) to facilitate systemic infection. Control animals received phosphate-buffered saline (PBS) via identical routes. The inoculation doses were selected based on prior ferret studies demonstrating reproducible systemic infection and to ensure sufficient viral dissemination for comparative evaluation of age- and genotype-dependent pathogenicity. FFU, focus-forming units; n, number of animals per group.

| Group | Treatment | Age(years) | Virus Strain | Virus titer (FFU/mL) | No. of animals |
| --- | --- | --- | --- | --- | --- |
| 1 | Infected | 3 | SFTSV B Type | 10<sup>7</sup> | 2 |
| 2 | Infected | 3 | SFTSV F Type | 10<sup>7</sup> | 2 |
| 3 | Control | 3 | PBS | - | 2 |
| 4 | Infected | 1 | SFTSV B Type | 10<sup>7</sup> | 2 |
| 5 | Infected | 1 | SFTSV F Type | 10<sup>7</sup> | 2 |
| 6 | Control | 1 | PBS | - | 2 |

Body weight and rectal temperature were measured daily from day 0 to day 7 post infection (dpi). Humane endpoints were predefined, and animals exhibiting ≥ 20% body weight loss or severe clinical deterioration were euthanized in accordance with ethical guidelines (KNU 2025-0836).

### Clinical scoring and sample collection

Clinical signs were assessed daily using a standardized scoring system based on activity, responsiveness, and physical appearance. In addition, rectal swab samples were collected daily from all animals to monitor viral shedding following SFTSV infection. Swabs were placed in viral transport medium and clarified by centrifugation prior to analysis. At the humane endpoint or scheduled necropsy, major organs including the liver, spleen, kidney, brain, small intestine, large intestine and blood were collected aseptically. Tissue samples were homogenized in sterile medium.

Viral loads in serum, rectal swabs, and tissue homogenates were quantified by real-time reverse transcription PCR kit (rRT-PCR, Powerchek SFTSV Real-time PCR Kit, Kogene Biothech, South Korea). A standard curve was generated using serial dilutions of SFTSV stocks with known infectious titers (FFU/mL), enabling conversion of viral loads quantified by rRT-PCR into FFU/mL equivalents.

### Hematological and biochemical analysis

Blood samples were collected at designated time points (0, 4, and 7 dpi) via venipuncture under light anesthesia. Whole blood was used for complete blood count (CBC) analysis, including white blood cell (WBC) and platelet (PLT) counts, while serum was separated for biochemical analysis Serum biochemical parameters, including aspartate aminotransferase (AST) and alanine aminotransferase (ALT), were measured using standard clinical chemistry assays according to the manufacturer’s instructions. These parameters were used to assess hematologic abnormalities and liver injury following SFTSV infection.

### Histopathological analysis

At the humane endpoint or scheduled necropsy, tissue samples from the liver and spleen were collected aseptically from all animals. Tissues were fixed in 4% paraformaldehyde(PFA) for at least 24 hours, processed using standard histological procedures, embedded in paraffin, and sectioned at a thickness of approximately 5 μm. Tissue sections were stained with hematoxylin and eosin (H&E) for histopathological evaluation.

### Ethics statement

All animal experiments were conducted in strict accordance with the guidelines for the care and use of laboratory animals and were approved by the Kyungpook National University Institutional Animal Care and Use Committee (IACUC) of the relevant institution (KNU 2025-0836). All procedures involving live SFTSV were performed in certified BSL-3 facilities (KCDC-21-3-02) by trained personnel. Every effort was made to minimize animal suffering, and humane endpoints were applied throughout the study.

### Statistical analysis

Data are presented as mean ± standard error of the mean (SEM). Statistical analyses were performed using GraphPad Prism software (version 5.0; GraphPad Software, San Diego, CA, USA). For experiments with sufficient sample sizes, comparisons between multiple groups were conducted using one-way or two-way analysis of variance (ANOVA), followed by appropriate post hoc tests. When the number of animals per group was limited (n = 2), data were analyzed descriptively without formal statistical testing. Differences were considered statistically significant at *p* < 0.05 where applicable.

## Acknowledgements

This study was supported by the Korea Disease and Prevention Agency (KCDA) and the National Institute of Health, Republic of Korea.

## Funding

This work was supported by the Korea Disease Control and Prevention Agency (KDCA) and the National Institute of Health, Republic of Korea under grant number 6634-332. The funders had no role in study design, data collection and analysis, decision to publish, or preparation of the manuscript.

## Conflicts of interest

The authors declare no conflicts of interest.

